## Supplementary material for "Rapid reconstruction of neural circuits using tissue expansion and lattice light sheet microscopy": Figures S1-S4

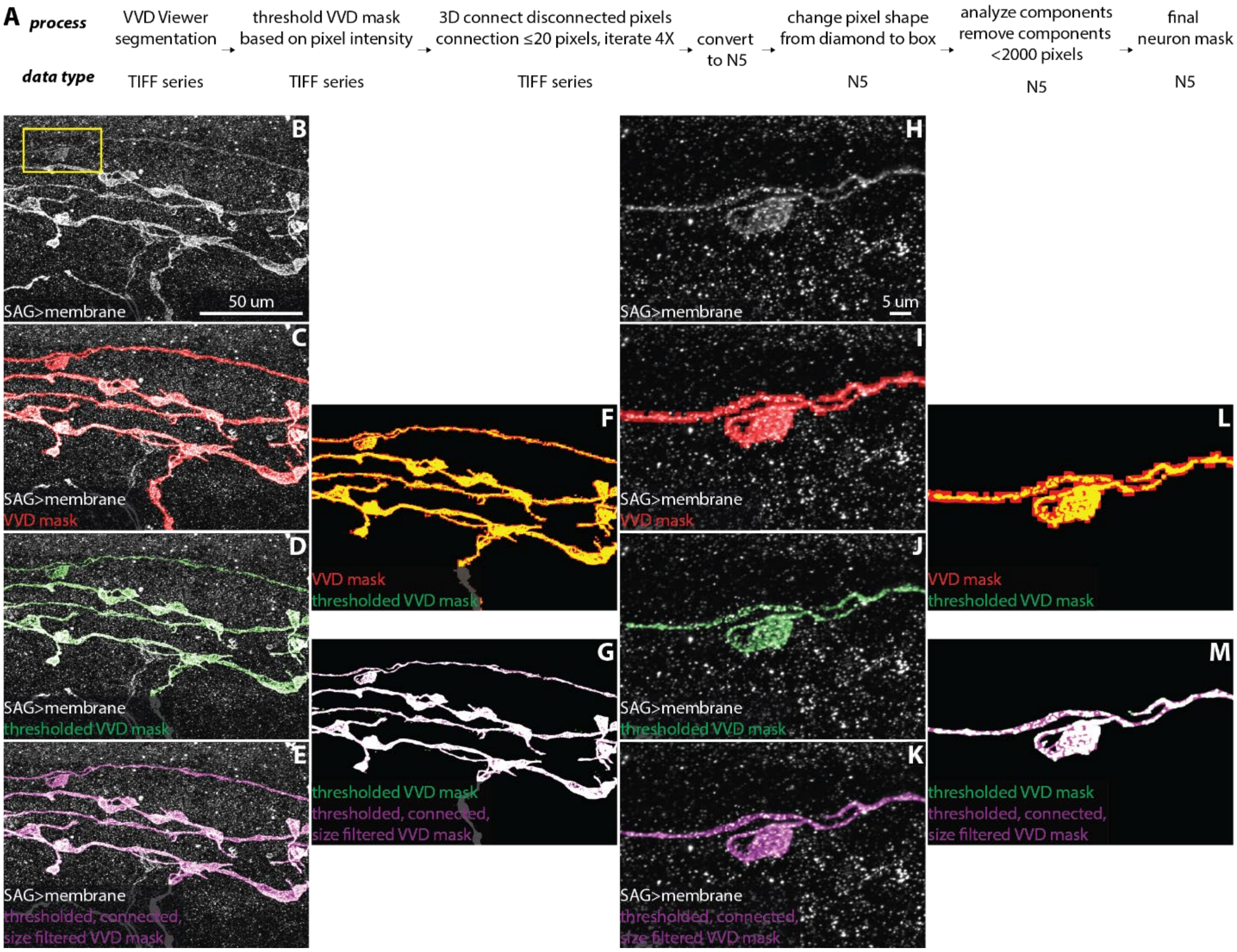


**Fig. S1.** VVD Viewer neuron segmentation and post-VVD mask processing. (A) Overview of VVD Viewer segmentation and automated mask processing pipeline. (B) A portion of an SAG neuron and off-target neuron in the region. (C) SAG neuron with VVD mask overlaid. (D) SAG neuron with the VVD mask after thresholding overlaid. (E) SAG neuron with the thresholded VVD mask after 3D gap filling, pixel shape change from diamond to box, and filtering to remove all objects smaller than 2000 pixels overlaid. (F) Overlay of VVD mask and VVD mask after thresholding. (G) Overlay of thresholded VVD mask and thresholded VVD mask after 3D gap filling and size filtering. (H-M) Zoom of region in the yellow box in (B) corresponding to panels (B-G), respectively.

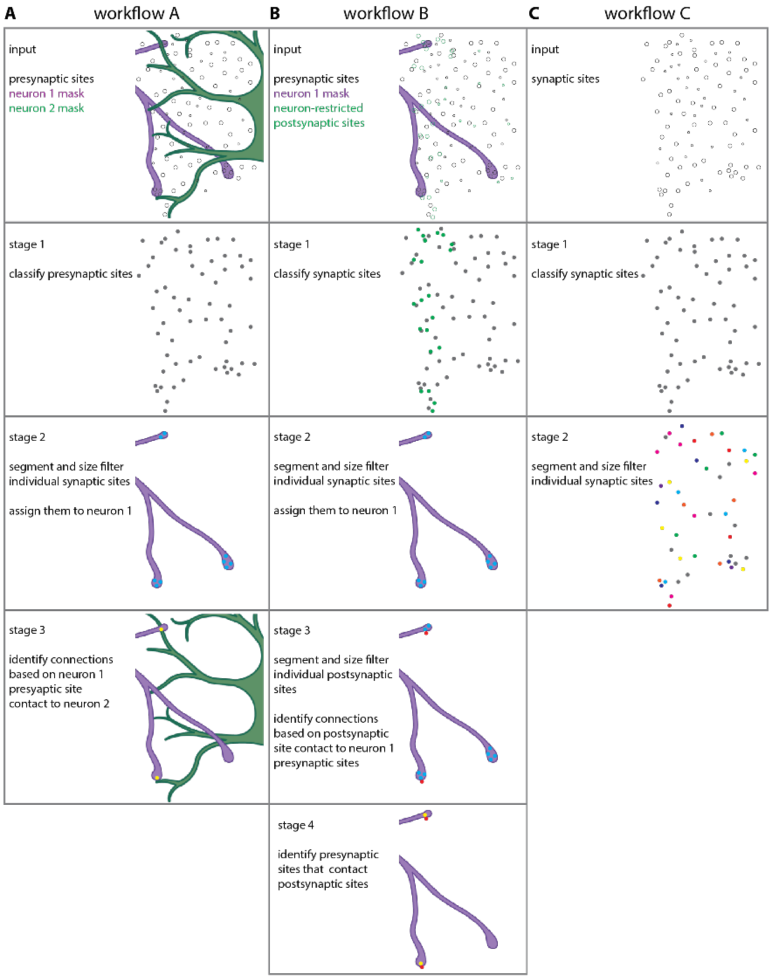


**Fig. S2.** Workflows for automatic synaptic site detection, assignment, and partner identification. (A) Workflow A quantifies presynaptic sites in neuron 1 and connections from neuron 1 to neuron 2 based on presynaptic site colocalization with the postsynaptic neuron (e.g. Fig. 3I-O, Fig. 5). (B) Workflow B quantifies presynaptic sites in neuron 1 and connections from neuron 1 to neuron 2 if presynaptic sites are labeled ubiquitously and postsynaptic sites are labeled specifically in neuron 2 (e.g. Fig. 3B-F). (C) Workflow C quantifies all pre- or postsynaptic sites in a given volume. (A-C) Stage 1 runs the trained U-Net model on synaptic sites (pre and/or post). (A-B) Stage 2 identifies individual presynaptic sites by running Watershed segmentation on the presynaptic site results from Stage 1 and a size filter to remove objects below a defined threshold. (A-B) Assigns the presynaptic sites to neuron 1 based on a defined colocalization threshold with the neuron 1 mask. (C) Stage 2 identifies individual synaptic sites by running Watershed segmentation on the synaptic site results from Stage 1 and a size filter to remove objects below a defined threshold. (A) Stage 3 identifies putative synaptic connections based on neuron 1 presynaptic sites identified in Stage 2 that colocalize with the neuron 2 mask. (B) Stage 3 identifies individual postsynaptic sites by running a Watershed segmentation on the postsynaptic site results from Stage 1 and a size filter to remove objects below a defined threshold. Connections are identified based on a colocalization between the postsynaptic sites and the neuron 1 presynaptic sites identified in Stage 2. (B) Stage 4 identifies the neuron 1 presynaptic sites that contact the postsynaptic sites that make connections identified in Stage 3.

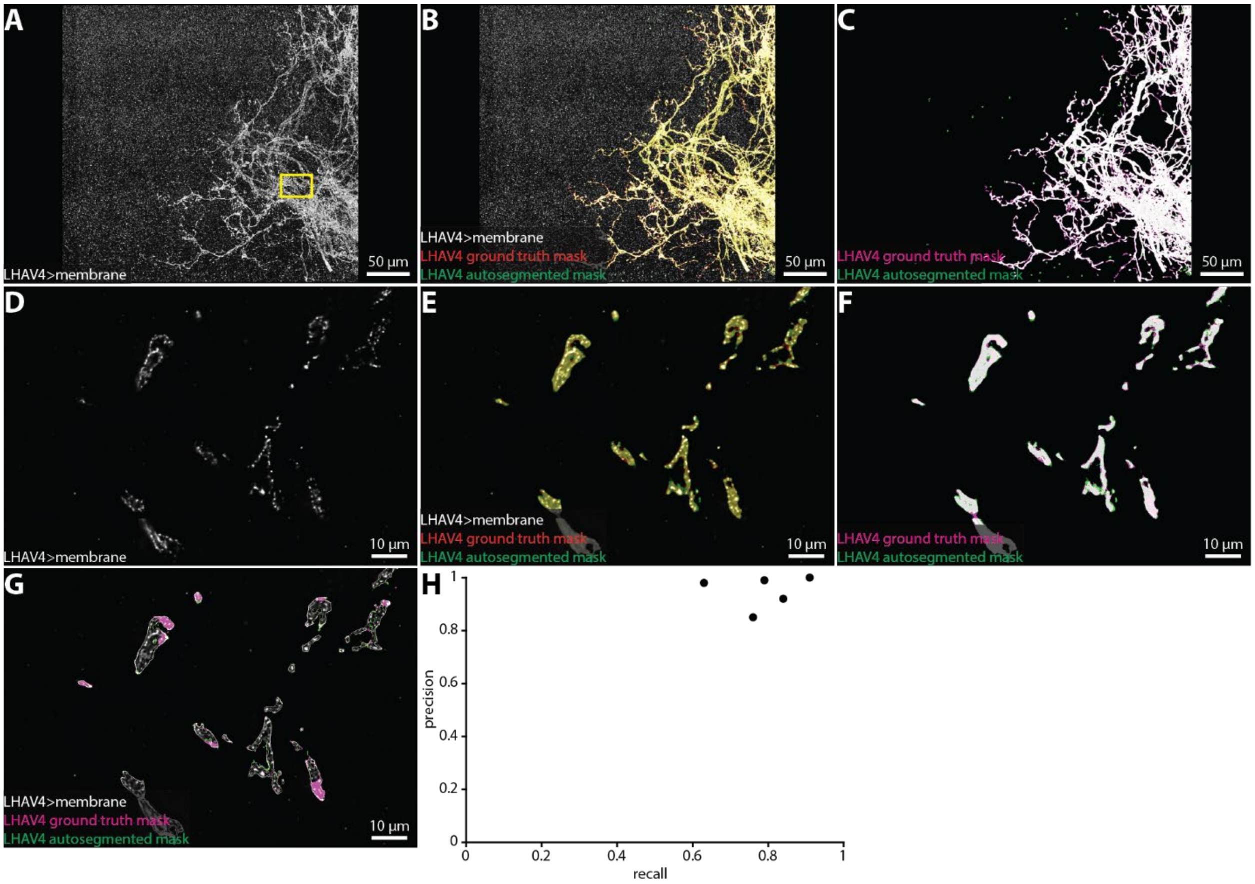


**Fig. S3.** Automatic neuron segmentation. Representative example of a group of neurons automatically segmented via a trained U-Net model. A subvolume of a group of lateral horn neurons (A, LHAV4) were segmented via the semi-automatic VVD Viewer segmentation workflow (LHAV4 ground truth mask) and via the U-Net automatic segmentation workflow (LHAV4 autosegmented mask) (B-C). (D-G) A single z-slice from the region in the rectangle in (A) corresponding to panels (B-C), respectively. (G) Overlay of LHAV4 neurons and neuron mask outlines. (H) Precision and recall plot of automatic neuron segmentation in five independent samples.

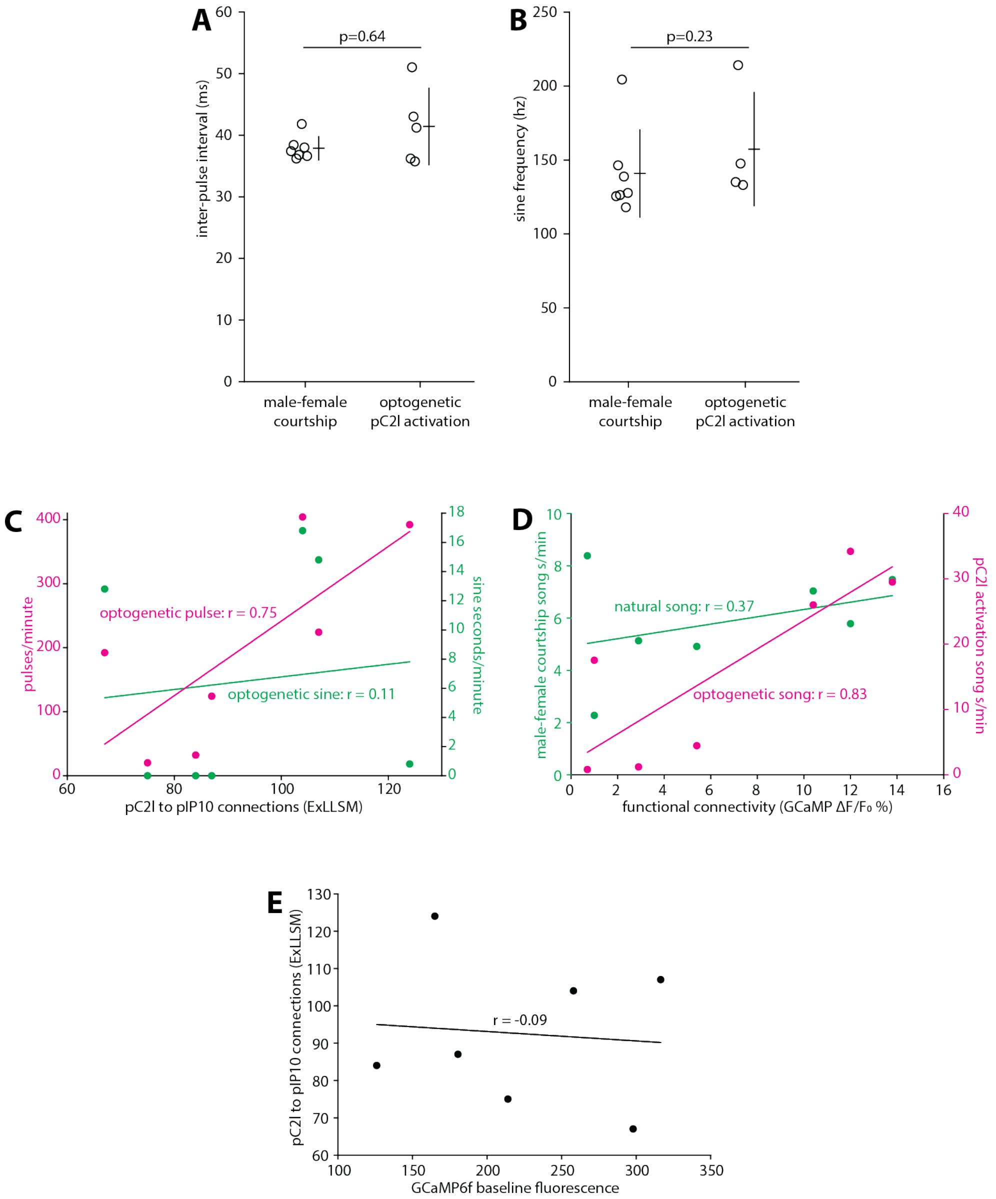


**Fig. S4.** (A) Inter-pulse interval of pulse trains produced during male-female courtship and during optogenetic pC2l activation (t-test, two-tailed P=0.64). For optogenetic pC2l activation, 2 flies produced isolated pulses without forming pulse trains. (B) Sine song frequency produced during male-female courtship and during optogenetic pC2l activation (t-test, two-tailed P=0.23). For optogenetic pC2l activation, 3 flies produced no sine song. (A-B) Individual animals and mean ± SD shown. (C) ExLLSM connections plotted against pulses/minute (magenta; extrapolated from 5 seconds of pC2l activation) and the sine seconds/minute (green; extrapolated from 5 seconds of pC2l activation) as in (Fig. 5B, H). (D) Functional pC2l>pIP10 connectivity plotted against song sec/minute produced during optogenetic pC2l activation (magenta; extrapolated from 5 seconds of pC2l activation) as in (Fig. 5B, H) and the song sec/minute produced during a 10-minute pairing with a female fly (green) as in (Fig. 5C, H). (E) Baseline GCaMP6f fluorescence (measured using identical parameters in each fly) plotted against the number of ExLLSM connections. (C-E) Linear regression and associated Pearson's Correlation Coefficient (r) values plotted for each relationship.
