## Supplementary material for "Rapid reconstruction of neural circuits using tissue expansion and lattice light sheet microscopy": Table 1

| Fig. | Genotype | Neurons>Reporters | Age | Sex | Primary Antibodies and Concentrations | Secondary Antibodies and Concentrations | Transgenic Fly Source |
|---|---|---|---|---|---|---|---|
| 1A | 10XUAS-IVS-myr::smGdP-HA (attP18), 13XLexAop2-IVS-myr::smGdP-FLAG (in su(Hw)attP8); VT000454-LexAGAD (attp40)/VT002064-p65ADZp; dsx-ZpGAL4DBD | SAG>smGdP-FLAG, pC1>smGdP-HA | 4 days old | F | Rabbit anti HA (1:200, Cell Signaling Technology, HA-Tag (C29F4) Rabbit mAb #3724); Rat anti FLAG (1:200, Novus Biologicals, DYKDDDDK Epitope Tag Antibody (L5) NBP1-06712); Mouse anti BRP (1:33, Developmental Studies Hybridoma Bank, nc82) | 1:500, Invitrogen, Goat anti-Rabbit IgG (H+L) Cross-Adsorbed Secondary Antibody, Alexa Fluor 568, A11011; 1:500, Invitrogen, Goat anti-Rat IgG (H+L) Cross-Adsorbed Secondary Antibody, Alexa Fluor 488, A11066; 1:500, Novus Biologicals, Goat anti-Mouse IgG (H+L) Secondary Antibody, Janelia Fluor 646, NBP1-72702JF646 | 10XUAS-IVS-myr::smGdP-HA, 13XLexAop2-IVS-myr::smGdP-FLAG (1); SAG (2); pC1 (3) |
| 1B-G | 10XUAS-IVS-myr::smGdP-HA (attP18), 13XLexAop2-IVS-myr::smGdP-FLAG (in su(Hw)attP8); VT044309-LexAGAD (attp40)/24C08-p65ADZp (attp40); UAS-Drep2-strawberry/VT043669-ZpGAL4DBD (attp2) | DA1-IPN>smGdP-FLAG, LHAV4>smGdP-HA (not shown); LHAV4>Drep2-strawberry (not shown) | 4-6 days old | M | Rabbit anti HA (1:200, Cell Signaling Technology, HA-Tag (C29F4) Rabbit mAb #3724); Rat anti FLAG (1:200, Novus Biologicals, DYKDDDDK Epitope Tag Antibody (L5) NBP1-06712); Mouse anti BRP (1:33, Developmental Studies Hybridoma Bank, nc82) | 1:500, Invitrogen, Goat anti-Rabbit IgG (H+L) Cross-Adsorbed Secondary Antibody, Alexa Fluor 568, A11011; 1:500, Invitrogen, Goat anti-Rat IgG (H+L) Cross-Adsorbed Secondary Antibody, Alexa Fluor 488, A11066; 1:500, Novus Biologicals, Goat anti-Mouse IgG (H+L) Secondary Antibody, Janelia Fluor 646, NBP1-72702JF646 | 10XUAS-IVS-myr::smGdP-HA, 13XLexAop2-IVS-myr::smGdP-FLAG (1); UAS-drep2-strawberry (4); DA1-IPN (5); LHAV4 (5) |
| 1H-P | 20XUAS-IVS-mCD8-GFP (attp5)/53G02-p65ADZp (attp40); 29G11-ZpGAL4DBD (attp2) | L2>membrane (not shown) | 6 days old | F | Chicken anti GFP (1:1000, Abcam, Anti-GFP antibody (ab13970)); Mouse anti BRP (1:33, Developmental Studies Hybridoma Bank, nc82) | 1:500, Invitrogen, Goat anti-Chicken IgY (H+L) Secondary Antibody, Alexa Fluor 488, A11039; 1:500 Goat anti-Mouse IgG (H+L) Highly Cross-Adsorbed Secondary Antibody, Alexa Fluor 568, A11031 | 20XUAS-IVS-mCD8-GFP (6); L2 (7) |
| 2B, D-G | 20XUAS-IVS-mCD8-GFP (attp5)/53G02-p65ADZp (attp40); 29G11-ZpGAL4DBD (attp2) | L2>membrane | 6 days old | F | Rabbit anti dsRed (1:1000, Takara Bio Living Colors® DsRed Polyclonal Antibody,632496); Mouse anti BRP (1:33, Developmental Studies Hybridoma Bank, nc82) | 1:500, Invitrogen, Goat anti-Rabbit IgG (H+L) Cross-Adsorbed Secondary Antibody, Alexa Fluor 568, A11011; 1:500, Invitrogen, Goat anti-Mouse IgG (H+L) Highly Cross-Adsorbed Secondary Antibody, Alexa Fluor 488, A11029 | 20XUAS-IVS-mCD8-GFP (6); L2 (7) |
| 2H-J | LexAop-myr-tdTomato/53G02-p65ADZp (attp40); UAS-FLP, brp-FRT-stop-FRT-V5-2A-LexA-VP16/29G11-ZpGAL4DBD (attp2) | L2>membrane, L2>STaR-BRP | 6 days old | F | Rabbit anti dsRed (1:1000, Takara Bio Living Colors® DsRed Polyclonal Antibody,632496); Mouse anti V5 (1:200, Mouse anti V5-Tag Antibody, clone SV5-Pk1, MCA1360) | 1:500, Invitrogen, Goat anti-Rabbit IgG (H+L) Cross-Adsorbed Secondary Antibody, Alexa Fluor 568, A11011; 1:500, Invitrogen, Goat anti-Mouse IgG (H+L) Highly Cross-Adsorbed Secondary Antibody, Alexa Fluor 488, A11029 | LexAop-myr-tdTomato; UAS-FLP, brp-FRT-stop-FRT-V5-2A-LexA-VP16 (8); L2 (7) |

| | | | | | | | |
|---|---|---|---|---|---|---|---|
| 2L, N-P | 10XUAS-IVS-myr::smGdP-HA (attP18), 13XLexAop2-IVS-myr::smGdP-FLAG (in su(Hw)attP8); VT044309-LexAGAD (attp40)/24C08-p65ADZp (attp40); UAS-Drep2-strawberry/VT043669-ZpGAL4DBD (attp2) | DA1-IPN>smGdP-FLAG, LHAV4>smGdP-HA (not shown); LHAV4>Drep2-strawberry (not shown) | 5 days old | F | Rabbit anti HA (1:200, Cell Signaling Technology, HA-Tag (C29F4) Rabbit mAb #3724); Rat anti FLAG (1:200, Novus Biologicals, DYKDDDDK Epitope Tag Antibody (L5) NBP1-06712); Mouse anti BRP (1:33, Developmental Studies Hybridoma Bank, nc82) | 1:500, Invitrogen, Goat anti-Rabbit IgG (H+L) Cross-Adsorbed Secondary Antibody, Alexa Fluor 568, A11011; 1:500, Invitrogen, Goat anti-Rat IgG (H+L) Cross-Adsorbed Secondary Antibody, Alexa Fluor 488, A11066; 1:500, Goat anti-Mouse IgG Secondary Antibody, ATTO 655 (STED), 15039 *DISCONTINUED | 10XUAS-IVS-myr::smGdP-HA, 13XLexAop2-IVS-myr::smGdP-FLAG (1); UAS-drep2-strawberry (4); DA1-IPN (5); LHAV4 (5) |
| 2R | 13XLexAop2-IVS-myr::smGFP-OLLAS [attP18]; VT00454LexAGAD (attp40)/VT002064-p65ADZp; 10XUAS-smFP-HA-drep2-sv40 (VK00005)/dsx-ZpGAL4DBD | SAG>smGdP-OLLAS, pC1>Drep2-HA (not shown) | 4 days old | F | Rat anti OLLAS (1:200, Novus Biologicals, OLLAS Epitope Tag Antibody (L2), NBP1-06713); Rabbit anti HA (1:200, Cell Signaling Technology, HA-Tag (C29F4) Rabbit mAb #3724); Mouse anti BRP (1:33, Developmental Studies Hybridoma Bank, nc82) | 1:500, Invitrogen, Goat anti-Rabbit IgG (H+L) Cross-Adsorbed Secondary Antibody, Alexa Fluor 568, A11011; 1:500, Invitrogen, Goat anti-Rat IgG (H+L) Cross-Adsorbed Secondary Antibody, Alexa Fluor 488, A11066; 1:500, Novus Biologicals, Goat anti-Mouse IgG (H+L) Secondary Antibody, Janelia Fluor 646, NBP1-72702JF646 | 13XLexAop2-IVS-myr::smGdP-OLLAS (1); 10XUAS-smFP-HA-drep2-sv40 (9); SAG (2); pC1 (3) |
| 3B-F | 13XLexAop2-IVS-myr::smGFP-OLLAS [attP18]; VT00454LexAGAD (attp40)/VT002064-p65ADZp; 10XUAS-smFP-HA-drep2-sv40 (VK00005)/dsx-ZpGAL4DBD | SAG>smGdP-OLLAS, pC1>Drep2-HA | 5 days old | F | Rat anti OLLAS (1:200, Novus Biologicals, OLLAS Epitope Tag Antibody (L2), NBP1-06713); Rabbit anti HA (1:200, Cell Signaling Technology, HA-Tag (C29F4) Rabbit mAb #3724); Mouse anti BRP (1:33, Developmental Studies Hybridoma Bank, nc82) | 1:500, Invitrogen, Goat anti-Rabbit IgG (H+L) Cross-Adsorbed Secondary Antibody, Alexa Fluor 568, A11011; 1:500, Invitrogen, Goat anti-Rat IgG (H+L) Cross-Adsorbed Secondary Antibody, Alexa Fluor 488, A11066; 1:500, Novus Biologicals, Goat anti-Mouse IgG (H+L) Secondary Antibody, Janelia Fluor 646, NBP1-72702JF646 | 13XLexAop2-IVS-myr::smGdP-OLLAS (1); 10XUAS-smFP-HA-drep2-sv40 (9); SAG (2); pC1 (3) |
| 3I-O | 10XUAS-IVS-myr::smGdP-HA (attP18), 13XLexAop2-IVS-myr::smGdP-FLAG (in su(Hw)attP8); VT000454-LexAGAD (attp40)/VT002064-p65ADZp; dsx-ZpGAL4DBD | SAG>smGdP-FLAG, pC1>smGdP-HA | 4 days old | F | Rabbit anti HA (1:200, Cell Signaling Technology, HA-Tag (C29F4) Rabbit mAb #3724); Rat anti FLAG (1:200, Novus Biologicals, DYKDDDDK Epitope Tag Antibody (L5) NBP1-06712); Mouse anti BRP (1:33, Developmental Studies Hybridoma Bank, nc82) | 1:500, Invitrogen, Goat anti-Rabbit IgG (H+L) Cross-Adsorbed Secondary Antibody, Alexa Fluor 568, A11011; 1:500, Invitrogen, Goat anti-Rat IgG (H+L) Cross-Adsorbed Secondary Antibody, Alexa Fluor 488, A11066; 1:500, Novus Biologicals, Goat anti-Mouse IgG (H+L) Secondary Antibody, Janelia Fluor 646, NBP1-72702JF646 | 10XUAS-IVS-myr::smGdP-HA, 13XLexAop2-IVS-myr::smGdP-FLAG (1); SAG (2); pC1 (3) |

| | | | | | | | |
|---|---|---|---|---|---|---|---|
| 4A-D | 10XUAS-IVS-myr::smGdP-HA (attP18), 13XLexAop2-IVS-myr::smGdP-FLAG (in su(Hw)attP8); 43924-LexAp65 (JK22C); 39575-GAL4 (attp2) | MB-DPM>smGdP-HA, MB-APL>smGdP-FLAG (not shown) | 4 days old | M | Rat anti HA (1:200, Roche, Anti-HA High Affinity from rat IgG1 11867423001); Mouse anti V5 (1:200, Mouse anti V5-Tag Antibody, clone SV5-Pk1, MCA1360); Rabbit anti INX6 (1:2500, Chia-Lin Wu) | 1:500, Invitrogen, Goat anti-Rabbit IgG (H+L) Highly Cross-Adsorbed Secondary Antibody, Alexa Fluor 488 A11029; 1:500, Invitrogen, Goat anti-Rat IgG (H+L) Cross-Adsorbed Secondary Antibody, Alexa Fluor 568 A11011; 1:500, Novus Biologicals, Goat anti-Mouse IgG (H+L) Secondary Antibody, Janelia Fluor 646, NBP1-72702JF646 | 10XUAS-IVS-myr::smGdP-HA, 13XLexAop2-IVS-myr::smGdP-FLAG (1); MB-DPM (10); MB-APL (11) |
| 4E-H | 10XUAS-IVS-myr::smGdP-HA (attP18), 13XLexAop2-IVS-myr::smGdP-FLAG (in su(Hw)attP8); 43924-LexAp65 (JK22C); 39575-GAL4 (attp2) | MB-DPM>smGdP-HA, MB-APL>smGdP-FLAG | 5 days old | F | Rat anti HA (1:200, Roche, Anti-HA High Affinity from rat IgG1 11867423001); Mouse anti V5 (1:200, Mouse anti V5-Tag Antibody, clone SV5-Pk1, MCA1360); Rabbit anti INX6 (1:2500, Chia-Lin Wu) | 1:500, Invitrogen, Goat anti-Rabbit IgG (H+L) Highly Cross-Adsorbed Secondary Antibody, Alexa Fluor 488 A11029; 1:500, Invitrogen, Goat anti-Rat IgG (H+L) Cross-Adsorbed Secondary Antibody, Alexa Fluor 568 A11011; 1:500, Novus Biologicals, Goat anti-Mouse IgG (H+L) Secondary Antibody, Janelia Fluor 646, NBP1-72702JF646 | 10XUAS-IVS-myr::smGdP-HA, 13XLexAop2-IVS-myr::smGdP-FLAG (1); MB-DPM (10); MB-APL (11) |
| 5E-F | 13XLexAop2-CsChrimson-tdTomato [attP18], 20XUAS-IVS-Syn21-opGCaMP6f-p10 [in su(Hw)attP8]; VT016273-p65ADZp (attP40), VT050279-ZpLexADBD (JK22C)/VT40556-p65ADZp; VT050279-ZpLexADBD (attP2)/VT40347-ZpGAL4DBD | pC2l>CsChrimson-tdTomato, pIP10>GCaMP6f | 5 days old | M | Rabbit anti dsRed (1:1000, Takara Bio Living Colors® DsRed Polyclonal Antibody,632496); Chicken anti GFP (1:1000, Abcam, Anti-GFP antibody (ab13970)); Mouse anti BRP (1:33, Developmental Studies Hybridoma Bank, nc82) | 1:500, Invitrogen, Goat anti-Rabbit IgG (H+L) Cross-Adsorbed Secondary Antibody, Alexa Fluor 568, A11011; 1:500, Invitrogen, Goat anti-Chicken IgY (H+L) Secondary Antibody, Alexa Fluor 488, A11039; 1:500, Novus Biologicals, Goat anti-Mouse IgG (H+L) Secondary Antibody, Janelia Fluor 646, NBP1-72702JF646 | 13XLexAop2-CsChrimson-tdTomato, 20XUAS-IVS-Syn21-opGCaMP6f-p10 (12); pIP10 (13); pC2l (5) |
| Sup. 1 | 10XUAS-IVS-myr::smGdP-HA (attP18), 13XLexAop2-IVS-myr::smGdP-FLAG (in su(Hw)attP8); VT000454-LexAGAD (attp40)/VT002064-p65ADZp; dsx-ZpGAL4DBD | SAG>smGdP-FLAG, pC1>smGdP-HA | 5 days old | F | Rabbit anti HA (1:200, Cell Signaling Technology, HA-Tag (C29F4) Rabbit mAb #3724); Rat anti FLAG (1:200, Novus Biologicals, DYKDDDDK Epitope Tag Antibody (L5) NBP1-06712); Mouse anti BRP (1:33, Developmental Studies Hybridoma Bank, nc82) | 1:500, Invitrogen, Goat anti-Rabbit IgG (H+L) Cross-Adsorbed Secondary Antibody, Alexa Fluor 568, A11011; 1:500, Invitrogen, Goat anti-Rat IgG (H+L) Cross-Adsorbed Secondary Antibody, Alexa Fluor 488, A11066; 1:500, Novus Biologicals, Goat anti-Mouse IgG (H+L) Secondary Antibody, Janelia Fluor 646, NBP1-72702JF646 | 10XUAS-IVS-myr::smGdP-HA, 13XLexAop2-IVS-myr::smGdP-FLAG (1); SAG (2); pC1 (3) |

| | | | | | | | |
|---|---|---|---|---|---|---|---|
| Sup. 3 | 10XUAS-IVS-myr::smGdP-HA (attP18), 13XLexAop2-IVS-myr::smGdP-FLAG (in su(Hw)attP8); VT044309-LexAGAD (attp40)/24C08-p65ADZp [attp40]; UAS-drep2-strawberry/VT043669-ZpGAL4DBD [attp2] | DA1-IPN>smGdP-FLAG (not shown), LHAV4>smGdP-HA, LHAV4>drep2-strawberry (not shown) | 4-6 days old | F | Rabbit anti HA (1:200, Cell Signaling Technology, HA-Tag (C29F4) Rabbit mAb #3724); Rat anti FLAG (1:200, Novus Biologicals, DYKDDDDK Epitope Tag Antibody (L5) NBP1-06712); Mouse anti BRP (1:33, Developmental Studies Hybridoma Bank, nc82) | 1:500, Invitrogen, Goat anti-Rabbit IgG (H+L) Cross-Adsorbed Secondary Antibody, Alexa Fluor 568, A11011; 1:500, Invitrogen, Goat anti-Rat IgG (H+L) Cross-Adsorbed Secondary Antibody, Alexa Fluor 488, A11066; 1:500, Novus Biologicals, Goat anti-Mouse IgG (H+L) Secondary Antibody, Janelia Fluor 646, NBP1-72702JF646 | 10XUAS-IVS-myr::smGdP-HA, 13XLexAop2-IVS-myr::smGdP-FLAG (1); DA1-IPN (5); LHAV4 (5) |

**Table 1:** Genotype, neurons, reporters, age, sex, antibodies, antibody concentrations, and transgenic fly source for data and figure panels.

**INX6 Antibody Source**

Wu CL, Shih MFM, Lai JSY, Yang HT, Turner GC, Chen L, Chiang AS (2011) Heterotypic Gap Junctions between Two Neurons in the Drosophila Brain Are Critical for Memory. Curr Biol 21:848–854.

**Transgenic Fly Source**

| | |
|---|---|
| 1 | Nern A, Pfeiffer BD, Rubin GM (2015) Optimized tools for multicolor stochastic labeling reveal diverse stereotyped cell arrangements in the fly visual system. Proc Natl Acad Sci U S A 112:E2967. |
| 2 | Feng K, Palfreyman MT, Häsemeyer M, Talsma A, Dickson BJ (2014) Ascending SAG Neurons Control Sexual Receptivity of Drosophila Females. Neuron 83:135-148. |
| 3 | Wang F, Wang K, Forknall N, Patrick C, Yang T, Parekh R, Bock D, Dickson BJ (2020) Neural circuitry linking mating and egg laying in Drosophila females. Nat 2020 5797797 579:101-105. |
| 4 | Andlauer TFM, Scholz-Kornehl S, Tian R, Kirchner M, Babikir HA, Depner H, Loll B, Quentin C, Gupta VK, Holt MG, Dipt S, Cressy M, Wahl MC, Fiala A, Selbach M, Schwärzel M, Sigrist SJ (2014) Drep-2 is a novel synaptic protein important for learning and memory. Elife 3:1-24. |
| 5 | This study |
| 6 | Pfeiffer BD, Ngo T-TB, Hibbard KL, Murphy C, Jenett A, Truman JW, Rubin GM (2010) Refinement of tools for targeted gene expression in Drosophila. Genetics 186:735-755. |
| 7 | Tuthill JC, Nern A, Holtz SL, Rubin GM, Reiser MB (2013) Contributions of the 12 neuron classes in the fly lamina to motion vision. Neuron 79:128. |
| 8 | Chen Y, Akin O, Nern A, Tsui CK, Pecot MY, Zipursky SL (2014) Cell-type Specific Labeling of Synapses in vivo through Synaptic Tagging with Recombination (STaR). Neuron 81:280. |
| 9 | Gift from Yoshi Aso |
| 10 | MB-DPM identified in this line by Yichun Shuai and Glenn Turner |
| 11 | Tanaka NK, Tanimoto H, Ito K (2008) Neuronal assemblies of the Drosophila mushroom body. J Comp Neurol 508:711-755. |

| | |
|---|---|
| 12 | Aso Y, Ray RP, Long X, Bushey D, Cichewicz K, Ngo TT, Sharp B, Christoforou C, Hu A, Lemire A, Tillberg P, Hirsh J, Litwin-Kumar A, Rubin GM (2019) Nitric oxide acts as a cotransmitter in a subset of dopaminergic neurons to diversify memory dynamics. Elife 8. |
| 13 | Ding Y, Lillvis JL, Cande J, Berman GJ, Arthur BJ, Long X, Xu M, Dickson BJ, Stern DL (2019) Neural Evolution of Context-Dependent Fly Song. Curr Biol 29:1089-1099.e7. |
